# Nanopore sequencing of the glucocerebrosidase (*GBA*) gene in a New Zealand Parkinson’s disease cohort

**DOI:** 10.1101/748335

**Authors:** O.E.E. Graham, T.L. Pitcher, Y. Liau, A.L. Miller, J.C. Dalrymple-Alford, T.J. Anderson, M.A. Kennedy

## Abstract

**Introduction:** Bi-allelic mutations in the gene for glucocerebrosidase (*GBA*) cause Gaucher disease, an autosomal recessive lysosomal storage disorder. Gaucher disease causing *GBA* mutations in the heterozygous state are also high risk factors for Parkinson’s disease (PD). *GBA* analysis is challenging due to a related pseudogene and structural variations (SVs) that can occur at this locus. We have applied and refined a recently developed nanopore DNA sequencing method to analyze *GBA* variants in a clinically assessed New Zealand longitudinal cohort of PD.

**Method:** We examined amplicons encompassing the coding region of *GBA* (8.9kb) from 229 PD cases and 50 healthy controls using the GridION nanopore sequencing platform, and Sanger validation.

**Results:** We detected 23 variants in 21 PD cases (9.2% of patients). We detected modest PD risk variant p.N409S (rs76763715) in one case, p.E365K (rs2230288) in 12 cases, and p.T408M (rs75548401) in seven cases, one of whom also had p.E365K. We additionally detected the possible risk variants p.R78C (rs146774384) in one case, p.D179H (rs147138516) in one case which occurred on the same haplotype as p.E365K, and one novel variant c.335C>T or p.(L335=), that potentially impacts splicing of *GBA* transcripts. Additionally, we found a higher prevalence of dementia among patients with *GBA* variants.

**Conclusion:** This work confirmed the utility of nanopore sequencing as a high-throughput method to identify known and novel *GBA* variants, and to assign precise haplotypes. Our observations may contribute to improved understanding of the effects of variants on disease pathogenesis, and to the development of more targeted treatments.

## Introduction

Several monogenic forms of Parkinson’s disease (PD) and many genetic risk factors, which predispose to development of PD, are now known. One of the strongest genetic risk factors for PD is mutation of the gene known as *GBA*, which encodes the lysosomal enzyme glucocerebrosidase (GCase) [1]. GCase is an enzyme involved in glycolipid metabolism, and mutations in *GBA* lead to a marked decrease in GCase activity [2]. Bi-allelic mutations in *GBA* result in excessive accumulation of the GCase substrate, GlcCer in macrophages, causing the rare lysosomal storage disorder, Gaucher disease (GD) [3, 4]. Heterozygous *GBA* mutations were initially believed to be benign; however, Parkinsonism was frequently reported in GD patients and their non-GD family members [5]. It is now clear that individuals with these *GBA* mutations associated with GD in the heterozygous state have a substantially elevated risk of developing PD compared with non-carriers [3, 6]. Furthermore, PD patients with *GBA* mutations have a noticeably earlier age of PD onset and a slightly increased risk of cognitive effects such as dementia, as well as a higher incidence of psychosis and delirium [7, 8].

Over 300 different *GBA* variants have been described, including single nucleotide polymorphisms (SNPs) (the majority of which are missense variants), frameshift mutations, splice-site alterations, and a range of structural variants (SVs) (i.e. variations >50bp) [4]. However, analysis of *GBA* is not straightforward. A nearby pseudogene called *GBAP1* shares 96% sequence homology with the coding region of the *GBA* gene, and care is required to prevent pseudogene contamination when analyzing *GBA* [9, 10].

Although application of a long-PCR step targeted at *GBA* can effectively solve many (but not all) of the issues arising from structural variants and the presence of a pseudogene [10], challenges remain using short read sequencing with such amplicons, including the inability to precisely assign haplotypes. Long read sequencing approaches such as Oxford Nanopore Sequencing (Oxford Nanopore Technologies; ONT) and PacBio (Pacific Biosciences) offer new opportunities for analyzing complex genes such as *GBA*. Indeed a recent report by Leija-Salazar *et al*. described a procedure for carrying out genotyping and haplotyping of *GBA* on the MinION sequencer (ONT) [11]. We have further refined the method of Leija-Salazar [11], and applied it for the discovery of *GBA* variants in a New Zealand PD longitudinal cohort. Our refined protocol utilizes updated hardware and software, thereby reducing the computational workload and improving sequencing accuracy.

## Materials and Methods

### Cohort description, recruitment and ethics

DNA for *GBA* genotyping was from a convenience sample (n = 229) of Parkinson’s disease patients from the New Zealand Brain Research Institute (NZBRI) PD cohort recruited through our movement disorders clinic for a range of PD-related research studies. We selected this convenience sample as our lab had previously collected blood samples from these patients for another study. Patients met the UK Brain Bank criteria for PD. Exclusion criteria included an indication of an atypical parkinsonian disorder, a history of moderate to severe head injury or stroke, a major psychiatric or medical illness in the previous six months, or poor English, which would prevent completion of neuropsychological testing. All participants had disease severity rated using the Movement Disorders Society Unified Parkinson’s Disease Rating Scale (MDS-UPDRS) and Hoehn and Yahr staging. Cognitive functioning was measured using a neuropsychological battery [12]. This analysis also used 50 age and sex-matched healthy controls recruited in conjunction with the PD patients. We collected blood samples from an arm vein into EDTA blood tubes. Blood was frozen at −20°C until DNA extraction. All participants provided informed consent. The Southern Health and Disability Ethics committee (New Zealand; URA/11/08/042 and URB/09/08/037) approved the study protocols.

### DNA preparation and PCR

DNA from peripheral blood samples was extracted using a modified salting-out method followed by a phenol-chloroform purification step [13]. For each sample, a PCR that generated an 8.9 kilobase (kb) amplicon encompassing the entire coding region of *GBA* was performed according to Leija-Salazar et al. [11]. Supplementary Table 1 lists the long PCR primers used.

For each sample, 100ng of genomic DNA was amplified in the presence of 1 unit of Kapa Hifi Hotstart ready mix and 0.3mM of each primer in a 50uL reaction. PCR consisted of 28 cycles of 95°C for 3 minutes, 98°C for 20 seconds, 60.9°C for 15 seconds, 72°C for 9 minutes, followed by a final extension at 72°C for 9 minutes.

#### Nanopore sequencing and validation of variants

We established the nanopore method using several positive control GD samples carrying known *GBA* mutations (purchased from Coriell Institute, USA). Coriell IDs for these samples were NA01607, NA00372, NA10873, NA20270, NA02627 and NA20273.

*GBA* amplicons were purified using High-prep magnetic beads, modified using a protocol to remove fragments < 3-4kb [14]. To attach barcodes for sample multiplexing, we adapted a method described in the LSK-109 protocol (ONT, Oxford, UK). Reactions consisted of 140ng of the first round PCR product, 1uL PCR barcode (PCR Barcoding Expansion 1-96, EXP-PBC096), and 1 unit of Kapa Hifi Hotstart Ready Mix in a 50uL reaction. Thermocycler conditions consisted of 28 cycles of 95°C for 3 minutes, 98°C for 20 seconds, 62°C for 15 seconds, 72°C for 9 minutes, and one final extension of 72°C for 9 minutes. The barcode purification, ligation of the nanopore adapters, library preparation, and flow cell priming was performed according to the LSK-109 protocol. Prepared libraries were run on R9.4 flowcells using the GridION platform (ONT) for between 24 and 48 hours with high-accuracy live base calling (Guppy v2.2.2 flip-flop).

#### Nanopore sequence data analysis

The software pipeline for processing nanopore FASTQ read files, and generating variant calls as VCF files was as follows. After filtering out reads with a quality score (q-score) < 7, we used Porechop v0.2.4 (https://github.com/rrwick/Porechop) to remove sequencing adapters and demultiplex the reads. Two different alignment tools, minimap2 v2.16 (r922) [15] and NGMLR v0.2.7 (https://github.com/philres/ngmlr), were used to map reads to the *GBA* reference sequence (derived from Hg38) and generate the BAM alignment files. We employed Qualimap v2.2 (http://qualimap.bioinfo.cipf.es/) and MinIONQC v1.4.1 (https://github.com/roblanf/minion_qc) to obtain quality control metrics for the alignments and the nanopore reads, respectively. As an additional quality control measure, read alignments to the human genome (Hg38) were visualized and manually inspected using Ribbon, a tool for visualizing complex genome alignments and structural variation [16].

SNPS and small insertions/ deletions (indels) were called as VCF files using Nanopolish v0.11.1 [17] using the standard settings. The parameter --fix-homopolymers was included to prevent indel calls in homo-polymeric regions, which are generally false positives.

To call SV’s (>50bp changes), we used the program Sniffles v1.0.11 (https://github.com/fritzsedlazeck/Sniffles) with the default parameters.

To exclude false positives, we divided the Nanopolish quality score for the variant call by the total number of reads associated with the call. Adjusted quality scores >1.8 were considered true variants and <1.2 were considered false positives, the same threshold used by Leija-Salazar et al. [11].

Alleles were phased using WhatsHap v0.18 [18] with the default parameters. The phased VCF files for each of the patients were merged into a single file, enabling variant assessment. File manipulation was performed using SAMtools v1.3.1 (https://github.com/samtools) and VCFtools v0.1.16 (https://vcftools.github.io). All variants were viewed on www.varsome.com, to assess their significance. Varsome provides a compilation of mutation data from gnomAD and dbSNP among others.

As a further measure to ensure that the variant reads were not a result of systematic error, we assessed a subset of the variant calls via Sanger sequencing (Supplementary Methods 1) and by manually inspecting the BAM files in IGV.

The haplotypes for each of the samples were extracted as FASTA files from the phased VCF file using the BCFtools v1.9 consensus parameter (http://samtools.github.io/bcftools/). Unique haplotypes were then determined using the R ‘haplotypes’ package (https://cran.r-project.org/web/packages/haplotypes), and aligned using MAFFT (https://mafft.cbrc.jp/alignment/software/). A median-joining haplotype network was constructed and visualized from the haplotype alignment using PopART [19].

The bioinformatics workflow is outlined in Figure 1.

**Figure 1.**
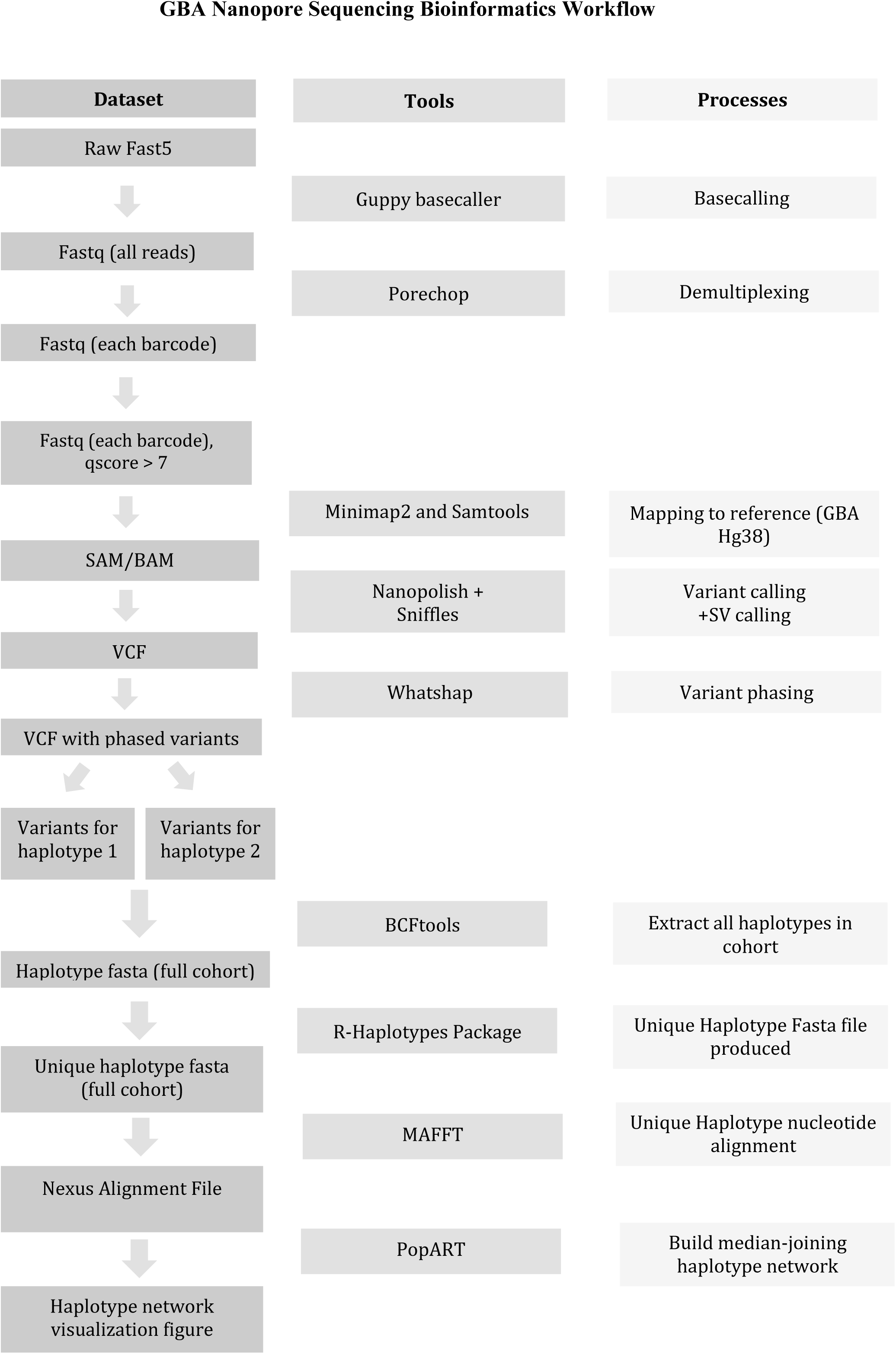
Bioinformatics workflow for *GBA* nanopore sequencing and haplotype analysis.

### Statistical Analysis

We calculated the significance of the relationship between *GBA* variant status and dementia using Fishers Exact Test. To test the association between *GBA* variant status and clinical features of PD we used unpaired T-tests. P-values ≤ 0.05 were defined as statistically significant.

## Results

### Nanopore sequencing of GBA in NZBRI cohort

We applied the nanopore sequencing method to examine *GBA* amplicons derived from 229 PD and 50 healthy control DNA samples from the NZBRI cohort. We sequenced the amplicons over five libraries containing multiplexed samples, on separate flow cells. All of the nanopore runs resulted in similar yields (~10-15 Gb with QC ≥ 7) (Supplementary Figure 1). The majority of these were the 8.9kb fragment of interest (Supplementary Figure 2). For the first run, ~60% of reads mapped to the *GBA* reference genome, and only a small fraction of the reads mapped to the pseudogene. Coverage depth over *GBA* averaged >2000x across all samples, far in excess of the 100x coverage required for accurate *GBA* genotyping [11].

We validated the efficacy of the protocol for *GBA* genotyping by successfully identifying and calling eight different mutations, including six SNPs, one indel and one recombination mutant in positive control GD samples (Coriell Institute, USA). The alignment program minimap2 was preferred for variant calling due to its speed, although we simultaneously used NGMLR for alignment prior to SV detection, due to its compatibility with the SV caller Sniffles.

By only considering variants with adjusted quality scores >1.8, all of the false positives were successfully filtered out, as verified by manually checking the BAM alignment files in Integrated Genomics Viewer (IGV) [20], and by Sanger sequencing validation of a subset of the predicted false positives.

### Potentially Pathogenic GBA variants discovered

Using this approach, out of the 229 NZBRI PD cohort samples, we identified six different variants of potential pathogenic significance over 21 different patients (Table 1) including two patients who each had two variants, and one patient with a homozygous variant (i.e. a total of 24 potential risk variants in 21 PD patients). We have denoted patients with these potential risk variants as *GBA* (*+*), and the other patients as *GBA* (-). We detected two patients out of 50 with potential risk variants in the healthy control population.

**Table 1.** Exon variants detected in this cohort

| **Amino acid change** | **Old variant nomenclature** | **Rs ID** | **Number of Samples** | **Number of Alleles** | **Cohort Allele Frequency** | **Control Population Allele Frequency** | **European non-Finnish Allele Frequency (GnomAD)** |
| --- | --- | --- | --- | --- | --- | --- | --- |
| p.E365K | p.E326K | rs2230288 | 12 | 12 | 0.0262009 | 0 | 0.01234 |
| p.T408M | p.T369M | rs75548401 | 7 | 8 | 0.0175439 | 0.020 | 0.009459 |
| p.D179H | p.D140H | rs147138516 | 1 | 1 | 0.0021930 | 0 | 0.000272 |
| p.N409S | p.N370S | rs76763715 | 1 | 1 | 0.0021930 | 0 | 0.002045 |
| c.335C>T;<br>p.(L335=) |  | Novel variant | 1 | 1 | 0.0021930 | 0 | N.A |
| p.R78C | p.R39C | rs146774384 | 1 | 1 | 0.0021930 | 0 | 0.000139 |
| **All Mutations** |  |  | **21\*** | 24 | 0.052517 | 0.020 |  |

\*Two of the patients carried two risk variants, and one patient was homozygous for one variant

The allele frequency of most of these variants in the cohort is elevated compared with the healthy control population, as well as the European non-Finnish population frequencies obtained from gnomAD (Table 1). The variant at the highest frequency in the cohort was p.E365K, with 12 carriers amongst cases. One patient in the cohort was heterozygous for both the p.E365K and the p.T408M variant. We identified one patient with the p.D179H variant who also carried p.E365K. In addition, one patient was found to have a novel variant c.335C>T; p.(L335=) at a splice selection site of exon 8 as indicated by Human Splice Finder (https://omictools.com/hsf-tool). We were unable to detect any SVs in the cohort. Additionally, we detected many variants of possibly benign or uncertain significance in the cohort, including a novel intron SNP (Supplementary Table 2).

In total, we detected 55 different unique haplotypes in the cohort, as visualized by a haplotype network analysis (Supplementary Figure 3).

### Clinical phenotypes of mutation carriers

In the PD cohort samples, there was a significantly (P < 0.05) higher prevalence of dementia (42.9%) in the *GBA* (+) group compared with the *GBA* (-) group (20.20%) (Table 2). Moreover, we noted a slightly decreased age at first symptom and diagnosis, albeit non-significant, in the *GBA* (+) carrier group (58.3 & 60.1 years, respectively), compared with the *GBA* (-) group (60.7 & 62.6 years). Age at PD dementia was also lower (71.3 years) in the *GBA* (*+*) group, along with disease duration until dementia (74.8 years), although these differences were non-significant.

**Table 2.**
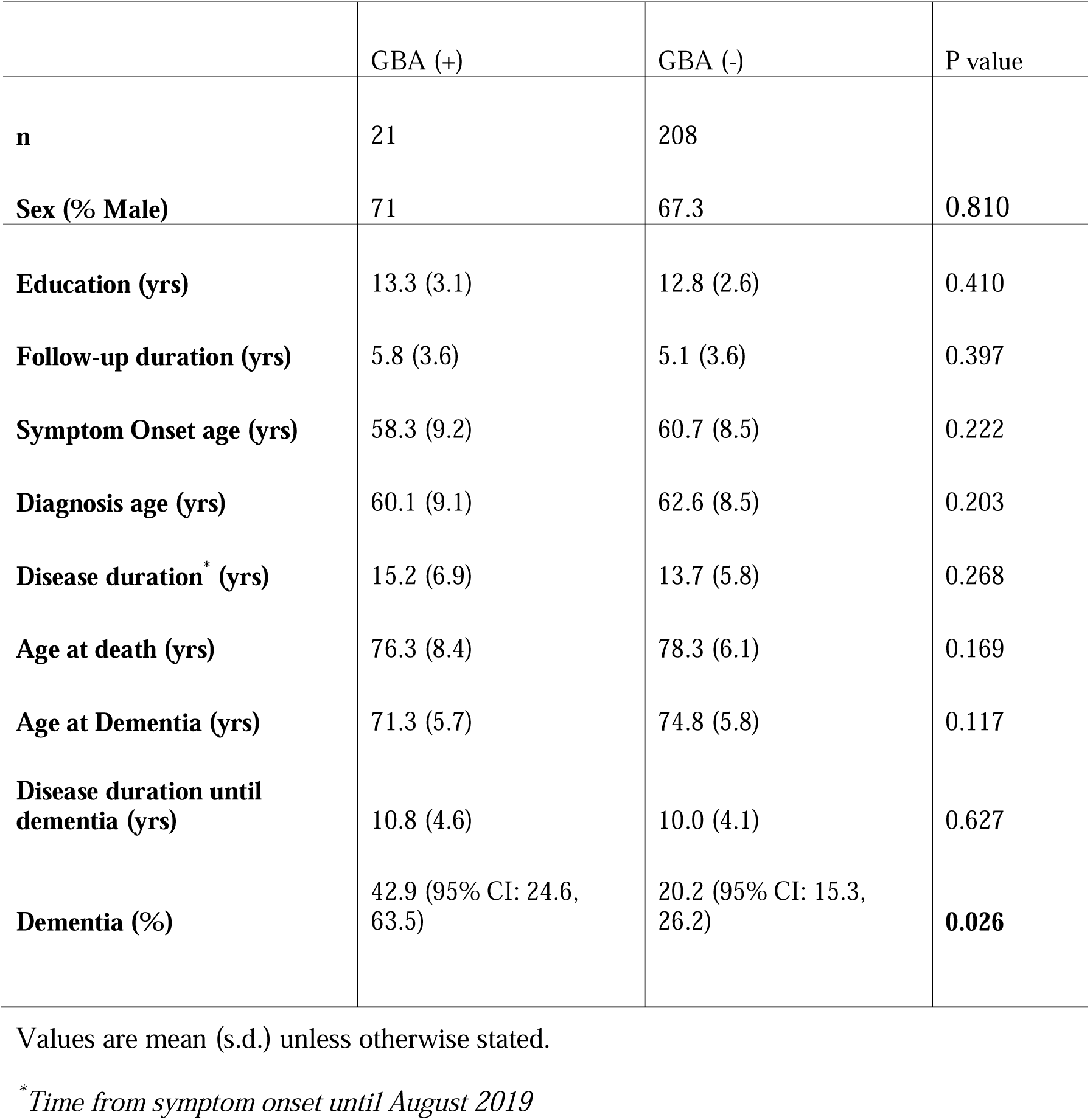
Demographic and clinical data for the PD patients in the NZBRI cohort separated by *GBA* mutation status.

|  | GBA (+) | GBA (-) | P value |
| --- | --- | --- | --- |
| **n** | 21 | 208 |  |
| **Sex (% Male)** | 71 | 67.3 | **0.810** |
| **Education (yrs)** | 13.3 (3.1) | 12.8 (2.6) | 0.410 |
| **Follow-up duration (yrs)** | 5.8 (3.6) | 5.1 (3.6) | 0.397 |
| **Symptom Onset age (yrs)** | 58.3 (9.2) | 60.7 (8.5) | 0.222 |
| **Diagnosis age (yrs)** | 60.1 (9.1) | 62.6 (8.5) | 0.203 |
| **Disease duration\* (yrs)** | 15.2 (6.9) | 13.7 (5.8) | 0.268 |
| **Age at death (yrs)** | 76.3 (8.4) | 78.3 (6.1) | 0.169 |
| **Age at Dementia (yrs)** | 71.3 (5.7) | 74.8 (5.8) | 0.117 |
| **Disease duration until dementia (yrs)** | 10.8 (4.6) | 10.0 (4.1) | 0.627 |
| **Dementia (%)** | 42.9 (95% CI: 24.6, 63.5) | 20.2 (95% CI: 15.3, 26.2) | **0.026** |

Values are mean (s.d.) unless otherwise stated.
\*Time from symptom onset until August 2019

## Discussion

### Nanopore sequencing as a tool for GBA mutation discovery

We successfully validated the protocol of Leija-Salazar et al. [11] as an effective method for genotyping of *GBA*. Additionally, we have updated the software pipeline utilized in the method of Leija-Salazar et al. [11], improving the accuracy and reducing the computational workload. More specifically our pipeline utilizes Guppy (flip-flop) instead of the now discontinued base-calling tool Albacore. Guppy runs orders of magnitude faster (1,500,000 bp/s vs 120,000 bps/s) due to its use of GPU acceleration [21]. The use of Guppy combined with the GridION device, which has a dedicated GPU for base-calling, enables fast, real-time base calling, a significant improvement over the CPU constrained Albacore. Another notable advantage of our protocol is the use of the read alignment tool Minimap2 instead of Graphmap. Minimap requires 3.6x less memory than Graphmap on average, and is 2.5x faster with no cost to overall accuracy [22].

Moreover, we highlight the capability of this protocol to phase variants, thereby enabling the establishment of direct molecular haplotypes. We noted that there are two main haplotypes and many rare ones (Supplementary Figure 3). Most of these rare haplotypes only differed from the main haplotypes by a single nucleotide. Understanding haplotypes may reveal useful information about the origins and inheritance of *GBA* variants.

We were unable to detect any structural variants (SVs) (variations >50bp), despite previous studies reporting such variants in PD patients [4, 23].

### Profile of GBA variants discovered in NZPD

In total, we detected variants potentially relevant to the context of PD in 9.2% of patients in the cohort, compared with 4.0% of patients in our healthy control cohort. This frequency is similar to what was anticipated based on previous studies [4].

#### Modest risk variants: p.N409S, p.E365K and p.T408M

Homozygous p.N409S mutations are the most common variants associated with GD. There is good evidence that p.N409S increases the risk for PD significantly [4, 6, 24]. The study by Blauwendraat [24] found that p.N409S is associated with a significant increased risk for PD (OR = 1.882, *P* = 0.002, 95% CI not provided).

The frequency of the p.E365K allele in our cohort is 0.0262, compared with the gnomAD frequency in the European (non-Finnish) population of 0.011. We did not detect any p.E365K alleles in our control cohort. All 11 cases with p.E365K were heterozygotes. p.E365K is not itself associated with GD, and only results in a mild loss-of-function enzyme. A recent meta-analysis indicates that there is a correlation between this mutation and PD in total populations (OR = 1.99; 95% CI = 1.57–2.51) [25]. That study also found that p.E365K predicted a more rapid progression of cognitive decline and motor symptom dysfunction.

The variant allele frequency of p.T408M in the cohort is 0.0175 compared with the gnomAD European (non-Finnish) frequency of 0.009459. However, the frequency was not elevated compared with the control population (0.02), as we detected two patients heterozygous for p.T408M. One of our six p.T408M positive patients is homozygous for this variant, the rest are heterozygotes. One patient also carried p.E365K on the other allele. p.T408M is not associated with GD, though it does result in decreased GCase enzyme activity [2]. A recent meta-analysis reported that p.T408M is associated with an increased risk for PD (OR = 1.78; 95% CI = 1.25–2.53) [26].

Interestingly, p.N409S, p.E365K and p.T408M are all frequently overrepresented in patients with REM-sleep behavior disorder (RBD). Patients with RBD have a significantly elevated risk of developing PD with dementia [3, 27, 28].

#### Other Possible Risk variants detected

p.R78C has an allele frequency of 0.000139 in the gnomAD European (non-Finnish) population. The clinical significance of this mutation remains unresolved, although it has a “pathogenic” computational verdict based on seven predictions from DANN, GERP, dbNSFP, FATHMM, MetaLR, MetaSVM, MutationAssessor and MutationTaster. However, there is no evaluation of this mutation in the context of GD or PD in the literature.

The gnomAD frequency for p.D179H is 0.00027 in European (non-Finnish) populations. The variant has been identified in GD patients [29], and UniProt classifies this variant as ‘disease’, but there are no studies regarding this mutation in a PD context. The patient carrying p.D179H also carries p.E365K, and allele phasing revealed that the two mutations exist on the same haplotype (Figure 2).

**Figure 2.**
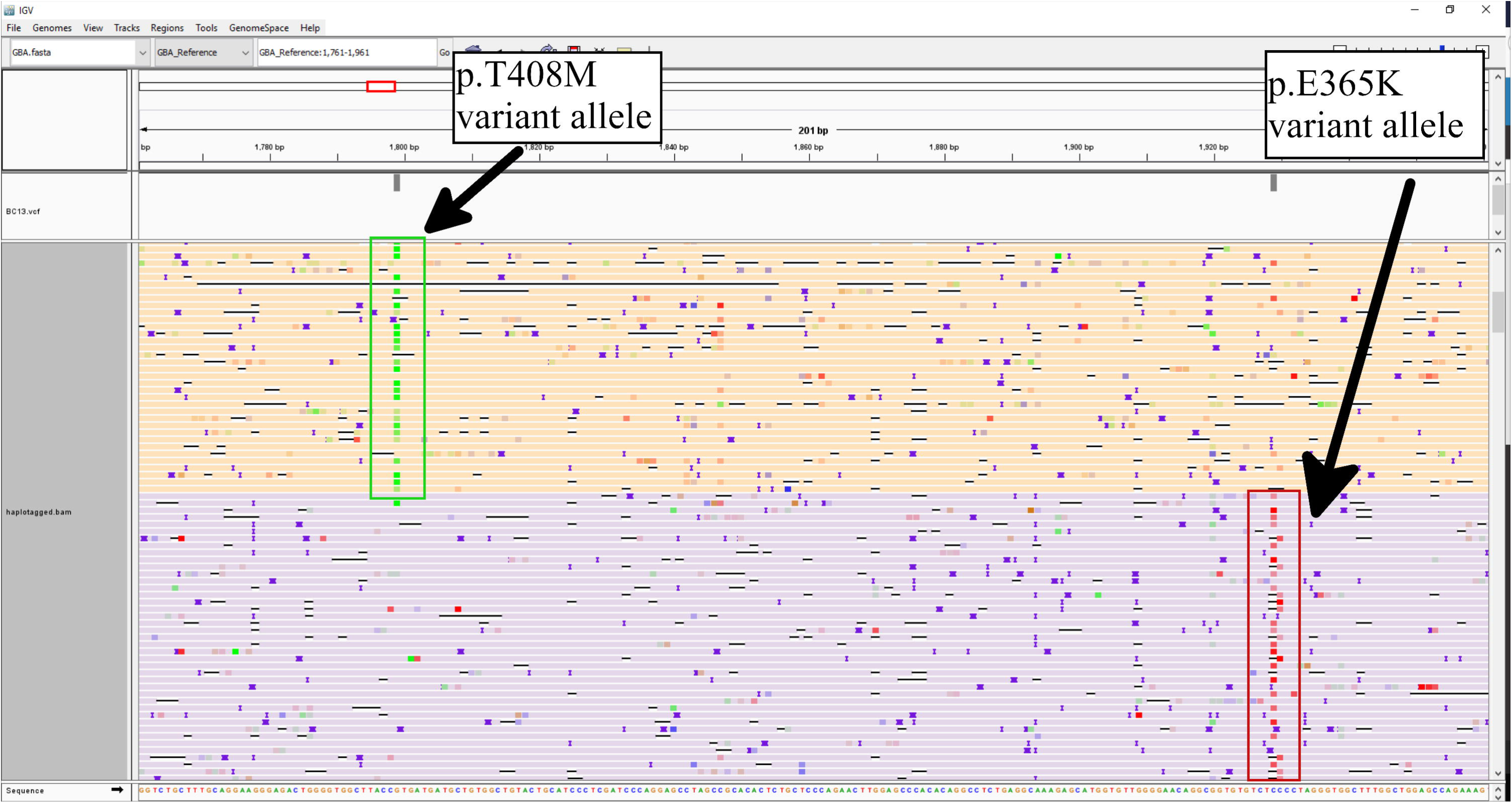
Example of *GBA* haplotypes generated by WhatsHap. Phased variants for the p.E365K/p.T408M double mutant as visualized in IGV. The different haplotypes are displayed in yellow (top) and purple (bottom). The top haplotype has the mutant allele at the p.T408M position (green box) and the ‘normal’ allele at the p.E365K position. The bottom haplotype has the mutant allele at the p.E365K position (red box) and the ‘normal’ allele at the p.T408M position.

We also detected c.335C>T; p.(L335=), a novel synonymous variant that it is predicted to affect splice enhancer function, although studies involving analysis of mRNA splicing patterns are needed to confirm this predicted function.

We were unable to detect any of these variants in the control cohort population. As we only have a single data point for each of the three risk variants, we are unable to make any claims regarding the role of these variants on clinical or developmental features of PD.

### Clinical implications

By comparing the clinical and demographic features of the patients between the *GBA* (+) and *GBA* (-) group, we provide support to previous studies indicating that PD patients with *GBA* mutations have a significantly higher prevalence of cognitive effects such as dementia [7]. Given the potential clinical effects of *GBA* mutations, and the interest in trialing drugs that target relevant pathways in *GBA* mutation positive patients (clinicaltrials.gov identifier: NCT02906020), application of the nanopore sequencing method to detect a wide range of mutations at high throughput will have considerable utility.

### Limitations

While we could not detect any SVs for a large portion of reads, sometimes we noted split reads between the gene and pseudogene in NGMLR alignments. These split reads could depict novel SVs, although they tended to appear in a small number of reads, meaning they are more likely to be chimeric PCR artefacts. Application of a novel PCR-free library generation method using CRISPR/ Cas9 capture of native DNA may be one possible solution to this limitation [30].

Another limitation of nanopore sequencing is its inability to accurately resolve homopolymers and detect small insertions and deletions in these regions. Given this limitation, we simply filtered out these variants, meaning we may have overlooked some true variants. Continual hardware and software improvements in this system are likely to overcome this problem in due course.

We also acknowledge that our cohort size is limited, preventing us from making statistical claims regarding the effect of *GBA* variants on clinical features of PD, or the allele frequency elevation in PD samples vs healthy controls. We addressed the issue of allele frequency elevation by also including European (non-Finnish) allele frequencies from gnomAD in our comparison, which are expected to be comparable to New Zealand European population allele frequencies.

## Conclusions

We have successfully validated ONT sequencing of *GBA* as an efficient and accurate method for detecting a wide range of variants in multiple patient samples, as well as resolving the haplotype of these variants. Applying this method to a sizeable New Zealand cohort of PD revealed the presence of multiple known *GBA* variants, and a range of additional variants of unclear clinical significance.

These findings may contribute to a better understanding of the effects of these variants on disease presentation or progression, and to development of more targeted treatments. This may be of importance as novel pharmaceuticals are actively under development to treat PD patients carrying *GBA* variants.

## Acknowledgements

The Health Research Council and Lotteries Health provided funding for sample collection and storage. We would also like to acknowledge NZBRI Assistant Research Fellows Leslie Livingston, Sophie Grenfell, and Bob Young for cognitive and clinical assessments, as well as NZBRI Research Fellows Dr Kyla Horne for cognitive testing, and Dr Michael MacAskill and Dr Daniel Myall for cohort database management.

## References

1. O’Regan, G., et al., Glucocerebrosidase Mutations in Parkinson Disease. J Parkinsons Dis, 2017. 7(3): p. 411–422.

2. Alcalay, R.N., et al., Glucocerebrosidase activity in Parkinson’s disease with and without GBA mutations. Brain: a journal of neurology, 2015. 138(Pt 9): p. 2648–2658.

3. Riboldi, G.M. and A.B. Di Fonzo, GBA, Gaucher Disease, and Parkinson’s Disease: From Genetic to Clinic to New Therapeutic Approaches. Cells, 2019. 8(4).

4. Sidransky, E. and G. Lopez, The link between the GBA gene and parkinsonism. The Lancet Neurology, 2012. 11(11): p. 986–998.

5. Halperin, A., D. Elstein, and A. Zimran, Increased incidence of Parkinson disease among relatives of patients with Gaucher disease. Blood Cells, Molecules, and Diseases, 2006.

6. Sidransky, E., et al., Multicenter analysis of glucocerebrosidase mutations in Parkinson’s disease. The New England journal of medicine, 2009. 361(17): p. 1651–1661.

7. Brockmann, K., et al., GBA-associated Parkinson’s disease: Reduced survival and more rapid progression in a prospective longitudinal study. Movement Disorders, 2015. 30(3): p. 407–411.

8. Oeda, T., et al., Impact of glucocerebrosidase mutations on motor and nonmotor complications in Parkinson’s disease. Neurobiology of Aging, 2015.

9. Tayebi, N., et al., Reciprocal and Nonreciprocal Recombination at the Glucocerebrosidase Gene Region: Implications for Complexity in Gaucher Disease, in Am J Hum Genet. 2003. p. 519–34.

10. Jeong, S.Y., et al., Identification of a novel recombinant mutation in Korean patients with Gaucher disease using a long-range PCR approach. J Hum Genet, 2011. 56(6): p. 469–71.

11. Leija-Salazar, M., et al., Evaluation of the detection of GBA missense mutations and other variants using the Oxford Nanopore MinION. Mol Genet Genomic Med, 2019. 7(3): p. e564.

12. Wood, K.-L., et al., Different PD-MCI criteria and risk of dementia in Parkinson’s disease: 4-year longitudinal study. Npj Parkinson&#39;S Disease, 2016. 2: p. 15027.

13. Miller, S.A., D.D. Dykes, and H.F. Polesky, A simple salting out procedure for extracting DNA from human nucleated cells. Nucleic acids research, 1988. 16(3): p. 1215–1215.

14. Nagar, R. and B. Schewessinger, DNA size selection (>3-4kb) and purification of DNA using an improved homemade SPRI beads solution. 2018.

15. Li, H., Minimap2: pairwise alignment for nucleotide sequences. Bioinformatics, 2018. 34(18): p. 3094–3100.

16. Nattestad, M., C.-S. Chin, and M.C. Schatz, Ribbon: Visualizing complex genome alignments and structural variation. bioRxiv, 2016: p. 082123.

17. Loman, N.J., J. Quick, and J.T. Simpson, A complete bacterial genome assembled de novo using only nanopore sequencing data. Nature Methods, 2015. 12: p. 733.

18. Martin, M., et al., WhatsHap: fast and accurate read-based phasing. bioRxiv, 2016: p. 085050.

19. Leigh, J.W. and D. Bryant, popart: full-feature software for haplotype network construction. Methods in Ecology and Evolution, 2015. 6(9): p. 1110–1116.

20. Robinson, J.T., et al., Integrative genomics viewer. Nature biotechnology, 2011. 29(1): p. 24–26.

21. Wick, R.R., L.M. Judd, and K.E. Holt, Performance of neural network basecalling tools for Oxford Nanopore sequencing. Genome Biology, 2019. 20(1): p. 129.

22. Senol Cali, D., et al., Nanopore sequencing technology and tools for genome assembly: computational analysis of the current state, bottlenecks and future directions. Briefings in Bioinformatics, 2018.

23. Spataro, N., et al., Detection of genomic rearrangements from targeted resequencing data in Parkinson’s disease patients. Movement Disorders, 2017. 32(1): p. 165–169.

24. Blauwendraat, C., et al., Coding variation in GBA explains the majority of the SYT11-GBA Parkinson’s disease GWAS locus. Movement disorders: official journal of the Movement Disorder Society, 2018. 33(11): p. 1821–1823.

25. Huang, Y., et al., The Association between E326K of GBA and the Risk of Parkinson’s Disease. Parkinson’s Disease, 2018. 2018: p. 6.

26. Mallett, V., et al., GBA p.T369M substitution in Parkinson disease: Polymorphism or association? A meta-analysis. Neurol Genet, 2016. 2(5): p. e104.

27. Iwaki, H., et al., Genetic risk of Parkinson disease and progression:: An analysis of 13 longitudinal cohorts. Neurology. Genetics, 2019. 5(4): p. e348–e348.

28. Gámez-Valero, A., et al., Glucocerebrosidase gene variants are accumulated in idiopathic REM sleep behavior disorder. Parkinsonism & related disorders, 2018. 50: p. 94–98.

29. Montfort, M., et al., Functional analysis of 13 GBA mutant alleles identified in Gaucher disease patients: Pathogenic changes and “modifier” polymorphisms. Hum Mutat, 2004. 23(6): p. 567–75.

30. Gilpatrick, T., et al., Targeted Nanopore Sequencing with Cas9 for studies of methylation, structural variants and mutations. bioRxiv, 2019: p. 604173.

