## Supplementary material for "Nanopore sequencing of the glucocerebrosidase (*GBA*) gene in a New Zealand Parkinson’s disease cohort": Supplementary Figure 2 Caption.docx

**Supplementary Figure 2.MinIONQC output showing the number of reads and the read lengths from a typical ONT run**. The top panel is all reads, the bottom panel is reads with at Q>7. The tall peak corresponds to the 8.9kb GBA amplicon, showing significant enrichment for GBA.
