## Supplementary material for "Nanopore sequencing of the glucocerebrosidase (*GBA*) gene in a New Zealand Parkinson’s disease cohort": Supplementary Figure 3 Caption.docx

**Supplementary Figure 3. Median-Joining Network of Haplotypes visualized using PopArt** [19]. Each node represents one of the 55 unique haplotypes within the cohort. The different nodes are colored according to the GBA variant present on the haplotype. The size of the node corresponds to the relative frequency of the haplotype within the PD cohort. Black nodes represent hypothetical haplotypes. Notches indicate 1bp differences. Numbers are internal haplotype ID codes.
