## Supplementary material for "Nanopore sequencing of the glucocerebrosidase (*GBA*) gene in a New Zealand Parkinson’s disease cohort": Supplementary Methods 1.docx

***Nested PCR:***

Nested PCRs covering different regions on the 1:1000 diluted 8.9kb *GBA* PCR product were performed. Nested PCRs were carried out in 10μL reactions containing 1μL of a 1:10,000 dilution of the long PCR product in the presence of 1x Fisher TaqTi Buffer, 0.4mM dNTPs, 1.5mM MgCl_2_, 0.05μL Fisher TAQ TI, 0.5mM of each primer and 1M betaine. Primers were designed using Geneious Prime® 2019.0.4 ([https://www.geneious.com](http://www.geneious.com/)) to enrich for exons containing the mutation of interest (Supplementary Table 1). DNA was amplified in a touchdown PCR consisting of 15 cycles of 94°C for 15 seconds, 70°C for 15 seconds, and 72°C for 1 minute, with annealing temperature decrease of 1°C per cycle. This was followed by 20 cycles of 94°C for 15 seconds, 55°C for 15 seconds, and 72°C for 1 minute and a final extension at 72°C for 5 minutes. Sanger sequencing was subsequently performed on the nested PCR products.

***Sanger Sequencing:***

Sanger sequencing was performed using 1 uL of 1:3 dilution of nested PCR product, 0.5mM primer (forward or reverse), 1x sequencing buffer, and 0.5uL of Big Dye Terminator (Thermo Fisher Scientific, Carlsbad, USA) in a 10uL reaction. Dye incorporation was performed in a thermo-cycler with 25 cycles of 96°C for 10 seconds, 50°C for 5 seconds, and 60°C for 4 minutes, and subsequently purified using Sephadex G-50 (Sigma-Aldrich, Saint Louis, USA). We utilized an AB3130XL Genetic Analyser (Thermo Fisher Scientific) to generate the Sanger traces. Sanger traces were visualised and converted to DNA sequences using Geneious Prime® 2019.0.4 ([https://www.geneious.com](http://www.geneious.com/)).

***Primer sequences:***

**Primers used for nested PCR amplification and Sanger Sequencing**

| **Targeted Region** | **Forward Primer** | **Reverse Primer** |
| --- | --- | --- |
| Exon4,5 | TTACTCTGTTGCCTGGGCTG | TCCCCTTCCTCCTCACCTTC |
| Exon6,7 | CTCAGAAAGGCCTGCGCTTC | GACCGAGAGAACAGGAAGCC |
| Exon9 | AGGTGTGCAGACCTGTGAAG | TGGGAGGATCAGTTGACCCT |
| Exon10 | CTGCCTCCATGGTGCAAAAG | TCTTCACACCCCCAACTCCT |
