## Supplementary material for "Nanopore sequencing of the glucocerebrosidase (*GBA*) gene in a New Zealand Parkinson’s disease cohort": Supplementary Table 1.docx

**Primers used for *GBA* Long PCR amplification including ONT index sequences**

| **Targeted Region** | **Forward Primer** | **Reverse Primer** |
| --- | --- | --- |
| *GBA (8.9kb)* | TCCTAAAGTTGTCACCCATACATG | CCAACCTTTCTTCCTTCTTCTCAA |
