## Supplementary material for "Nanopore sequencing of the glucocerebrosidase (*GBA*) gene in a New Zealand Parkinson’s disease cohort": Supplementary Table 2.docx

**Supplementary Table 2: *GBA* variants detected in the 229 Patients from the NZBRI PD cohort**

| **RS ID** | **Clinical Significance** | **Type** | **Reference Allele Frequency** | **Alternative Allele Frequency** |
| --- | --- | --- | --- | --- |
| **rs375776699** | Uncertain Significance | 3' UTR | A:0.971491 | G:0.0285088 |
| **rs368275143** | Uncertain Significance | 3' UTR | A:0.986842 | G:0.0131579 |
| **rs708606** | Uncertain Significance | 3' UTR | C:0.991228 | T:0.00877193 |
| **rs2974924** | Uncertain Significance | Intron | A:0.989035 | G:0.0109649 |
| **rs1800473** | Likely benign | Intron | T:0.713974 | C:0.286026 |
| **rs12752133** | Uncertain Significance | Intron | C:0.973799 | T:0.0262009 |
| **rs76763715** | PD Risk Allele | Missense | T:0.997807 | C:0.00219298 |
| **rs3115534** | Likely benign | Intron | G:0 | T:1 |
| **rs1335297441** | Uncertain Significance | intron | A:0.997807 | C:0.00219298 |
| **rs75548401** | Moderate PD Risk | Missense | G:0.982456 | A:0.0175439 |
| **rs2230288** | Moderate PD Risk | Missense | C:0.973799 | T:0.0262009 |
| **Novel (possible ESE variant)** | Possible Risk Variant | Synonymous | C:0.997807 | T:0.00219298 |
| **rs183903019** | Uncertain Significance | Intron | G:0.997807 | A:0.00219298 |
| **rs9628662** | Likely benign | Intron | T:0.713974 | G:0.286026 |
| **rs760954677** | Uncertain Significance | Intron | C:0.997807 | CA:0.00219298 |
| **rs72704130** | Uncertain Significance | Intron | G:0.964912 | A:0.0350877 |
| **rs140335079** | Uncertain Significance | Intron | A:0.97807 | T:0.0219298 |
| **rs762488** | Likely benign | Intron | T:0.716157 | C:0.283843 |
| **rs2009578** | Likely benign | Intron | G:0.716157 | A:0.283843 |
| **rs28678003** | Likely benign | Intron | C:0.964912 | T:0.0350877 |
| **Novel variant** | Uncertain Significance | Intron | G:0.997807 | A:0.00219298 |
| **rs183540501** | Uncertain Significance | Intron | G:0.995614 | T:0.00438596 |
| **rs2974923** | Likely benign | Intron | T:0.71179 | C:0.28821 |
| **rs147138516** | Possible Risk Variant | Missense | C:0.997807 | G:0.00219298 |
| **rs188328778** | Uncertain Significance | Intron | A:0.995614 | G:0.00438596 |
| **rs7416991** | Likely benign | Intron | T:0.165939 | C:0.626638 G:0.207424 |
| **rs556071493** | Uncertain Significance | Intron | G:0.997807 | A:0.00219298 |
| **rs2075569** | Likely benign | Intron | C:0.713974 | T:0.286026 |
| **rs146774384** | Possible Risk Variant | Missense | G:0.997807 | A:0.00219298 |
| **rs114217696** | Uncertain Significance | Intron | C:0.997807 | T:0.00219298 |
| **rs546949267** | Uncertain Significance | Intron | G:0.997807 | T:0.00219298 |
| **rs2361534** | Uncertain Significance | Intron | T:0.995614 | C:0.00438596 |
| **rs188978150** | Uncertain Significance | Intron | T:0.997807 | C:0.00219298 |

*Varsome (www.varsome.com) was used to assess the clinical significance of each variant*
