## Supplementary figures and images for "Nanopore sequencing of the glucocerebrosidase (*GBA*) gene in a New Zealand Parkinson’s disease cohort"

### Supplementary Figure 1.png

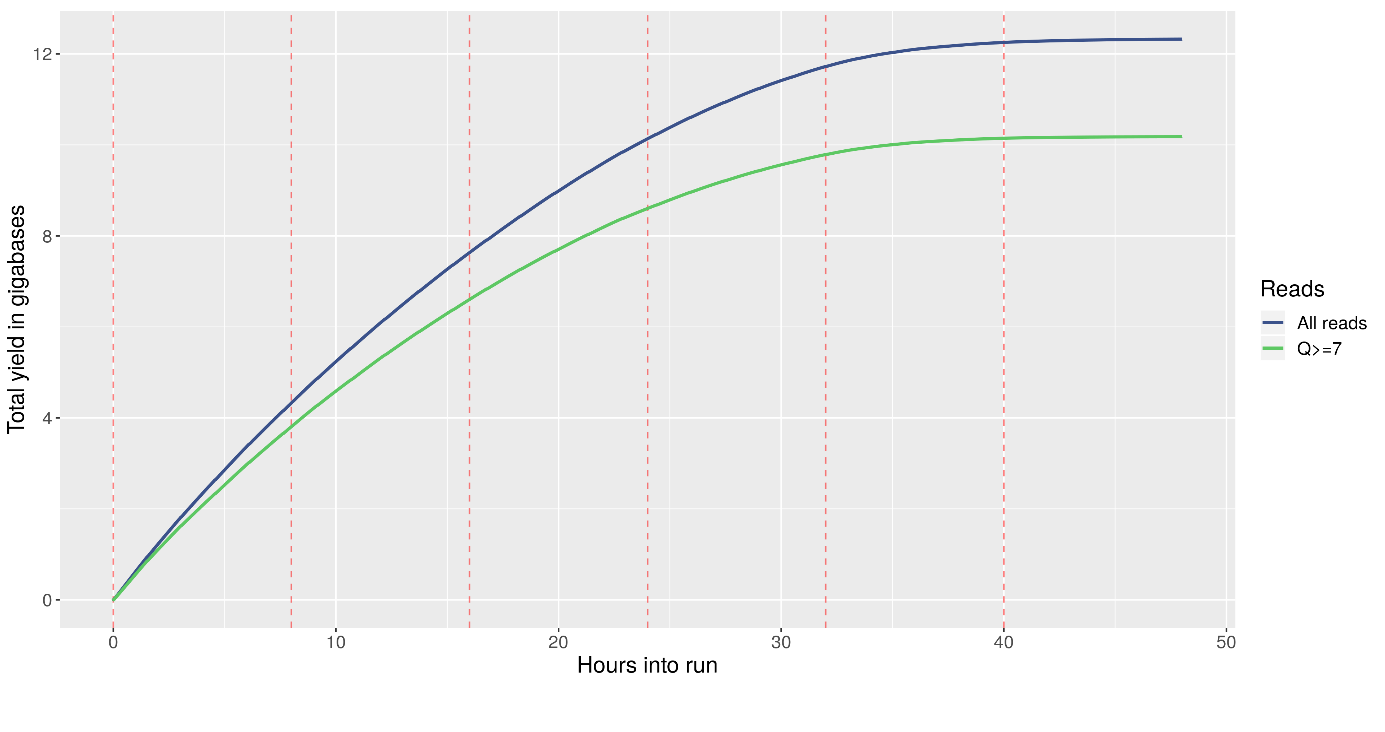

### Supplementary Figure 2.png

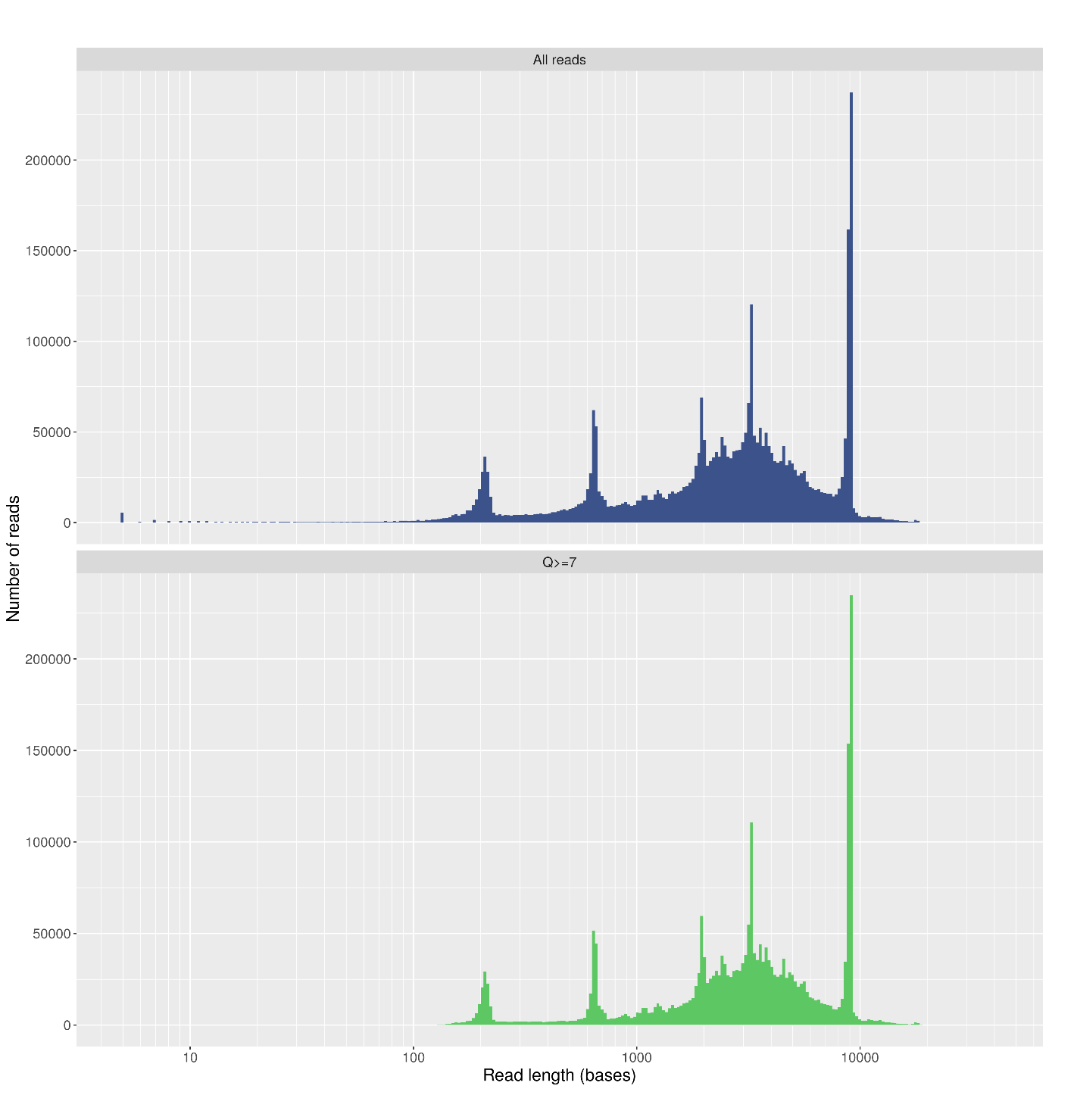

### Supplementary Figure 3.png

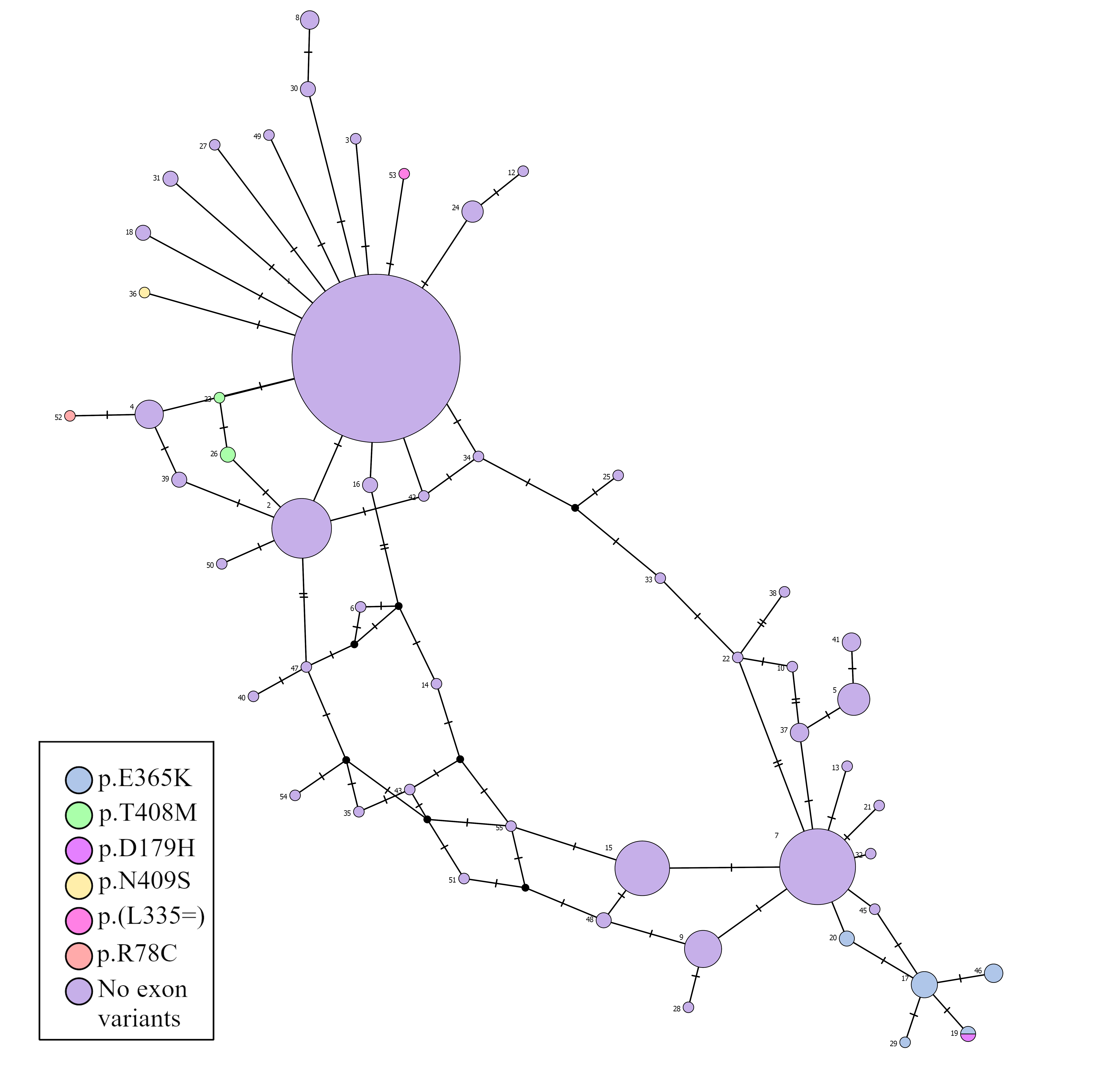
